# Matrix Metalloproteinase-2 as a novel regulator of glucose utilization by adipocytes

**DOI:** 10.1101/2024.12.09.626845

**Authors:** Melissa D. Lempicki, Ryan J. Garrigues, Tonya N. Zeczycki, Brandon L. Garcia, Thurl E. Harris, Akshaya K. Meher

## Abstract

Glucose transporter 4 (GLUT4) expression on white adipocytes is critical for absorbing excess blood glucose, failure of which promotes hyperglycemia. Matrix metalloproteinases (MMPs) play a crucial role in remodeling the white adipose tissue (WAT) during obesity. MMPs have multiple protein substrates, and surprisingly, it is unknown if they can directly target GLUT4 on the adipocyte surface and impair glucose absorption. We identified MMP2 as the highly active gelatinase, a class of MMP, in the gonadal WAT of high-fat diet-induced obese mice. *In vitro*, metabolic studies in 3T3-L1 adipocytes revealed MMP2 attenuated glucose absorption and glycolysis, which were recovered by an MMP2 inhibitor. *In silico* structural analysis using AlphaFold identified a putative MMP2 cleavage site on the extracellular domain of GLUT4. Further, in a substrate competition assay, a peptide mimicking the MMP2 cleavage motif on GLUT4 attenuated the cleavage of an MMP substrate by MMP2. Altogether, our results suggest a novel mechanism of impaired glucose absorption by adipocytes, which may contribute to hyperglycemia during obesity.

**ARTICLE HIGHLIGHTS:**

- It is unknown if MMPs directly affect adipocyte metabolism *via* targeting adipocyte surface proteins.
- We aimed to determine the highly active gelatinases, a class of MMP, in the white adipose tissue of obese mice, and examined its effect on adipocyte glucose metabolism.
- MMP2 was the highly active gelatinase in obese adipose tissue and, *in vitro*, MMP2 reduced adipocyte glucose metabolism, in part, by targeting the GLUT4 receptor.
- A harmful role of MMP2 is implicated in cardiometabolic diseases and our novel finding on the role of MMP2 on glucose metabolism may add to the pathogenesis of the diseases.

## INTRODUCTION

Obesity and its related comorbidity insulin resistance (IR), pose a significant health risk in developed countries. While many factors contribute to the development of IR, murine models of high-fat diet (HFD)-induced obesity revealed that glucose transporter 4 (GLUT4) expression on white adipocytes promotes insulin-stimulated glucose absorption and maintenance of whole-body glucose homeostasis (1). During obesity, GLUT4 expression on the surface of adipocytes decreases, leading to hyperglycemia and eventually IR (2). Matrix metalloproteinases (MMPs), a family of over 23 zinc-dependent endoproteinases, play a crucial role in remodeling the extracellular matrix of the white adipose tissue (WAT) during obesity (3). While MMPs are crucial for matrix remodeling, it is unknown whether MMPs directly affect adipocyte function and metabolism by degrading GLUT4 on the adipocyte surface.

Here, we identified MMP2 as the highly active gelatinase, a class of MMP, in the gonadal WAT of obese mice and determined if MMP2 directly affects adipocyte glucose utilization using 3T3-L1 adipocytes.

## RESEARCH DESIGN AND METHODS

The ‘Online Supplemental Materials’ provides details of the ‘Research Design and Methods’.

### Mice

Eight-week-old male C57BL/6J mice (#380050, Jackson Laboratory, Bar Harbor, ME) were given water and fed a high-fat diet (HFD, #F3282, Bio-Serv; 60% of calories from fat) for five to sixteen weeks or with a normal chow diet (NCD, Prolab IsoPro RMH 3000 5P76, LabDiet) for 11 or 16 weeks. All protocols involving animals were approved by the East Carolina University Animal Care and Use Committee and performed in an AAALAC-accredited facility in accordance with current NIH guidelines.

### Glucose Uptake

3T3-L1 adipocytes, 5 days after differentiation, were used for the glucose uptake assays using a fluorescent glucose analog 2-NBDG. Adipocytes were stimulated with insulin (200 nM) and treated at the same time with activated MMP2 (200, 400, or 800 ng/mL) for 20 min at 37 °C. For the MMP2 inhibitor assay, 1 μM MMP2 inhibitor was added to insulin and MMP2 (400 ng/ml)-treated adipocytes. Cytochalasin D (10 μM) was used as a negative control for insulin-stimulated glucose uptake.

### Seahorse

The 3T3-L1 fibroblasts on XFe24 well microplates coated with a poly-L-lysine solution and differentiated into adipocytes as described earlier (4) or seeded one day prior to seahorse experiments in poly-L-lysine coated microplates. After various treatments, a glycolysis stress test was performed using a Seahorse XFe24 Extracellular Flux Analyzer as described previously (5).

### MMP Substrate Cleavage Assay

In black 96-well plates, 5 nM (400 ng/mL) of MMP2 (BioLegend, #554402), 1 μM of MMP2 inhibitor (Cayman Chemicals, #19644), 5 nM of MMP2 with 1 μM of MMP2 inhibitor, or 5 nM of MMP2 with 50 mM EDTA as negative control were incubated with 5 μM of fluorescent ‘520 MMP FRET Substrate’ (AnaSpec) for 30 minutes in MMP substrate buffer. MMP activity was measured as at 494/521 excitation/emission.

For the substrate competition assay, 5 nM of MMP2 with 500 μM of GLUT4 loop peptide (RQGPGGPDSIPQGTL, GenScript), 500 μM of negative control peptide (LTGQPISDPGGPGQR, GenScript), 500 μM positive control peptide (VPLSLYSG, Bachm, #4109381), or 50 mM EDTA were incubated with 5 μM of ‘520 MMP FRET Substrate’ for 30 minutes in MMP substrate buffer.

### Statistical Analysis

The data were analyzed using GraphPad Prism 8 (GraphPad Software, La Jolla, CA) and presented as means ± SEM. Statistical analyses are provided in each figure legend. Differences between the groups were considered significant when P < 0.05.

## RESULTS

### MMP2 is the highly active gelatinase in white adipose tissue during obesity

In obese and IR humans and mice, macrophages infiltrate the adipose tissue and produce pro-inflammatory cytokines that can exacerbate IR development (6). Furthermore, plasma MMP levels correlate with several obesity-related parameters (7). To determine the highly active gelatinase in the WAT during HFD-induced obesity, we placed male C57BL6/J mice on an HFD for 5, 9, or 11 weeks or NCD for 11 weeks. As expected, the mice on HFD had increased body weight, increased fasting insulin levels, and reduced AKT phosphorylation in the gonadal WAT (**Sup. Fig. 1A-C)**. Apart from this, mice 11 weeks on HFD had an increase in the percent of total macrophages and an increase in expression of CD86, a marker for pro-inflammatory M1 macrophages, and an increase in the *Tnfα* gene expression in the stromal vascular fraction of the gonadal WAT (**Sup. Fig. 1D-F**). Furthermore, the number of crown-like structures (CLSs) was increased in the gonadal WAT at 5 weeks after HFD feeding (**Fig. 1A**). Interestingly, staining the adipose tissue sections with an MMP substrate, which becomes fluorescent upon cleavage, revealed increased MMP activity after HFD feeding, and the activity was primarily localized in the CLSs (**Fig. 1B**).

**Figure 1.**
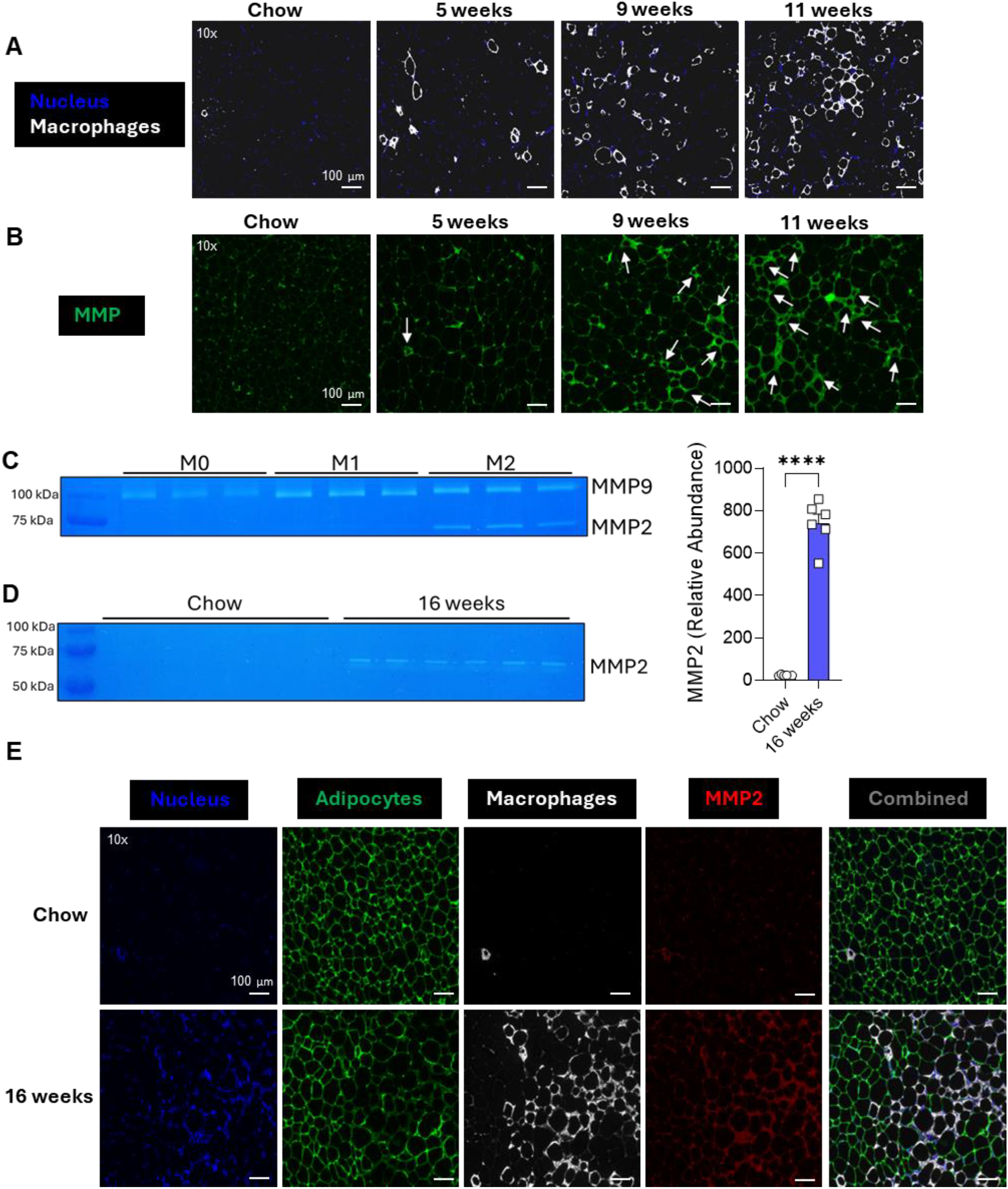
MMP2 is the highly active MMP in white adipose tissue during obesity. **(A)** Representative confocal images of gonadal WAT from mice on HFD for 5, 9, or 11 weeks or NCD for 11 weeks stained for nucleus (DAPI-blue), macrophages (Mac2-white). **(B)** Representative images of gonadal WAT from HFD for 5, 9, or 11 weeks or NCD for 11 weeks stained with fluorescent MMP substrate (green). **(C)** Gelatin zymogram of culture supernatant from M0 (unpolarized), M1 (pro-inflammatory), M2 (anti-inflammatory). **(D)** Gelatin zymogram of gonadal WAT lysate from mice on HFD or NCD for 16 weeks. **(E)** Representative images of gonadal WAT from mice on HFD or NCD for 16 weeks stained for nucleus (DAPI-blue), adipocytes (FABP4-green), macrophages (Mac2-white), MMP2 (red). Values are expressed as means + SEM. ****, p<0.0001 by parametric unpaired t-test. n=4 mice, 1 section per mouse, 3-4 images per section **(A-B)**, n=3 **(C)**, n=6 **(D)**, n=6 mice, 1 section per mouse, 3-4 images per section **(E)**. Scale bars: 100 μm **(A, B, & E)**

CLSs within the adipose tissue are comprised of macrophages that adopt a metabolically activated phenotype characterized by increased lysosomal activity and increased expression of pro-inflammatory cytokines, comparable to M1 activation (8). While pro-inflammatory cytokines have been shown to increase MMP expression, it is unclear what type of gelatinase are produced by M1-like pro-inflammatory macrophages, such as those found in the CLSs (9). We polarized BMDMs to M1 (pro-inflammatory) or M2 (anti-inflammatory) and analyzed the MMPs produced in the supernatant with a gelatin zymogram. Interestingly, M0 (unpolarized), M1, and M2 macrophages all produced MMP9 while only M2 macrophages produced MMP2 (**Fig. 1C**). To confirm the *in vitro* results, we examined MMP activity in the gonadal WAT lysate from mice on an NCD or HFD for 16 weeks and surprisingly, only MMP2 activity was detected in the adipose tissue after HFD feeding (**Fig. 1D**). To determine if other gelatinases were active in the earlier time points of HFD feeding, we performed zymography of gonadal WAT lysate of mice on HFD for 5 to 11 weeks and found only MMP2 activity which appeared as early as 5 weeks after HFD (**Sup. Fig. 1G)**. Immunohistochemistry staining of gonadal WAT showed increased MMP2 expression around the CLSs in the HFD mice compared to the NCD controls **(Fig. 1E**). These results suggest that MMP2 is the highly active gelatinase present in the gonadal WAT that localizes to CLSs after induction of HFD-induced obesity.

### MMP2 decreases glucose uptake and glycolysis in 3T3-L1 adipocytes

MMP2 has been previously shown to be secreted by the adipose tissue, and elevated MMP2 plasma levels can be found in obese patients; however, it remains unclear if MMP2 is having effects within the adipose tissue unrelated to its role in extracellular matrix remodeling (10). Therefore, we treated 3T3-L1 adipocytes with varying physiologically relevant concentrations of MMP2 and performed a glycolysis stress test to measure aerobic glycolysis using a Seahorse XFe24 Extracellular Flux Analyzer. Treatment with 400 or 800 ng/mL of MMP2 significantly decreased aerobic glycolysis, glycolytic capacity, and glycolytic reserve (**Fig. 2A**). Furthermore, treatment with an MMP2 inhibitor, which blocks MMP2’s catalytic domain (**Sup. Fig. 2A**), rescued aerobic glycolysis in 3T3-L1 adipocytes (**Fig. 2B**). To determine if decreased glycolysis was because of decreased glucose absorption, we examined glucose uptake in 3T3-L1 adipocytes after treatment with varying concentrations of MMP2 (**Sup. Fig. 2B**). In agreement with the results from the glycolysis stress test, glucose uptake was significantly decreased after treatment with 400 or 800ng/mL of MMP2 (**Fig. 2C & Sup. Fig. 2C**) and reduced glucose uptake was reversed after treatment with the MMP2 inhibitor (**Fig. 2D & Sup. Fig. 2D**). Altogether, these results indicate that MMP2, *via* protease activity, impairs aerobic glycolysis in adipocytes by decreasing glucose uptake into the cell.

**Figure 2.**
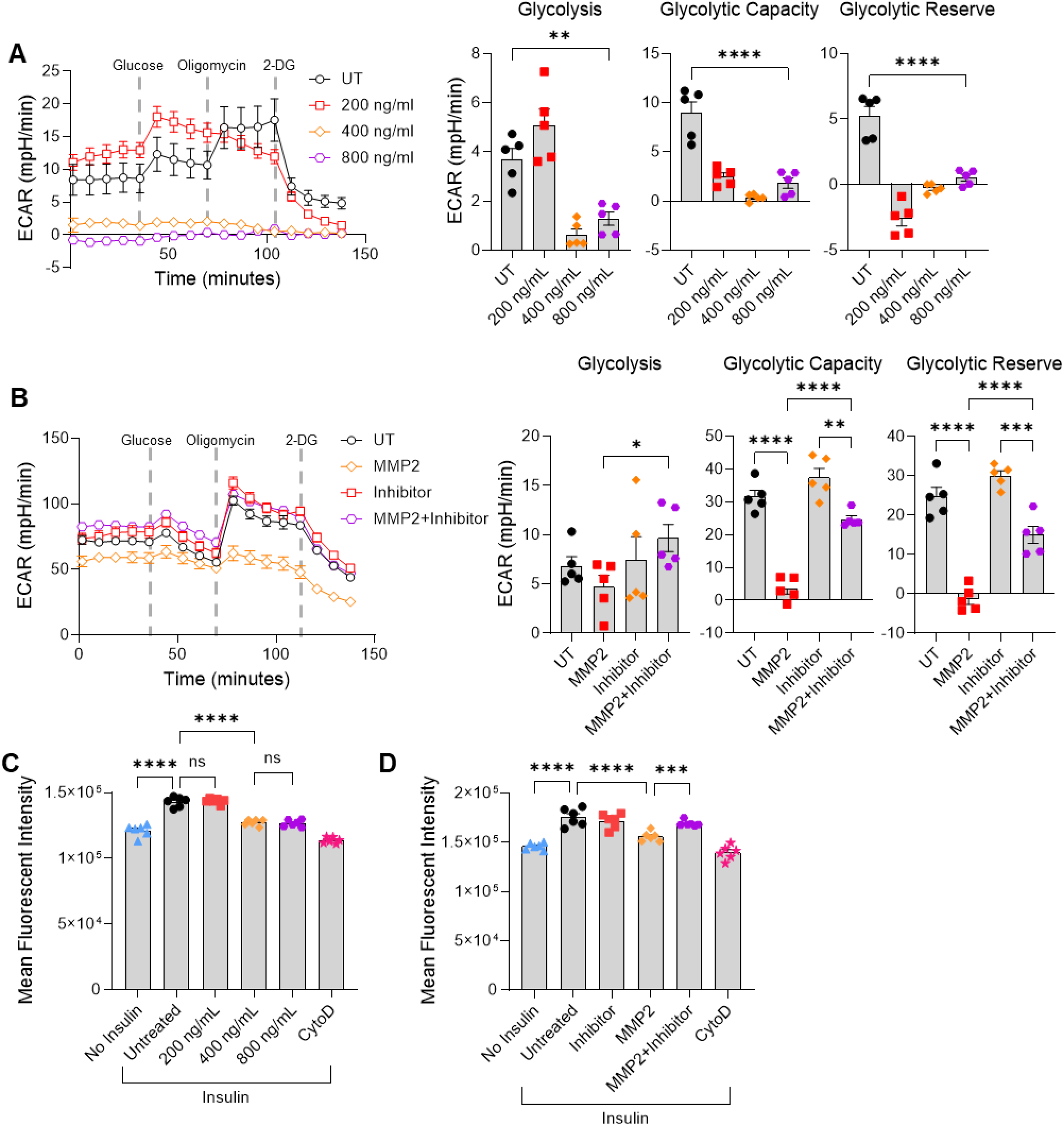
MMP2 decreases glucose uptake and glycolysis in 3T3L1 adipocytes. **(A)** Aerobic glycolysis of adipocytes after treatment with 200 ng/mL, 400 ng/mL, or 800 ng/mL of active MMP2. ECAR, extracellular acidification rate. **(B)** Aerobic glycolysis of adipocytes after treatment with 400 ng/mL of active MMP2, MMP2 inhibitor, or MMP2 with MMP2 inhibitor. **(C)** Glucose uptake of 3T3-L1 adipocytes after treatment with 200 ng/mL, 400 ng/mL, or 800 ng/mL of active MMP2. Glucose uptake was assessed by mean fluorescent intensity of 2-NBDG in live adipocytes. **(D)** Glucose uptake of 3T3-L1 adipocytes after treatment with 400 ng/mL of active MMP2, MMP2 inhibitor, or MMP2 with MMP2 inhibitor. Glucose uptake was assessed by mean fluorescent intensity of 2-NBDG in live adipocytes. Values are expressed as means + SEM. *, p<0.05; **, p<0.01; ***, p>0.001, ****, p<0.0001 by parametric unpaired t-test. n=5 **(A & B**) n=6 **(C & D)**

### The extracellular domain of the GLUT4 receptor contains a putative MMP2 cleavage site

Insulin-dependent glucose influx by 3T3-L1 adipocytes largely relies on cell surface GLUT4 (11). Amino acid sequence analysis of GLUT-4 identified an extracellular loop between the first and second helices that contains a putative MMP2 cleavage motif PXXXHy (12). XHy represents a large hydrophobic residue in the P1’ position; in GLUT4 it is isoleucine. AlphaFold2 (AF2) *in silico* folding analysis, utilizing the identified murine GLUT4 extracellular loop (Uniprot: P14142, residues 64-77) and full-length murine MMP2 (Uniprot: P33434), conformed the peptide to the active site of MMP2 in a substrate-like fashion with isoleucine occupying the P1’ pocket adjacent to the catalytic glutamine (**Fig. 3A**). To verify the *in silico* prediction, we created a GLUT4 loop peptide (residues 63-77) and performed an enzymatic competition assay with an MMP2 substrate which becomes fluorescent upon cleavage. Addition of the GLUT4 loop peptide significantly decreased the fluorescent signal indicating the GLUT4 loop peptide is, at least, binding to the substrate binding site of MMP2 and reducing its ability to cleave the substrate (**Fig. 3B**).

**Figure 3.**
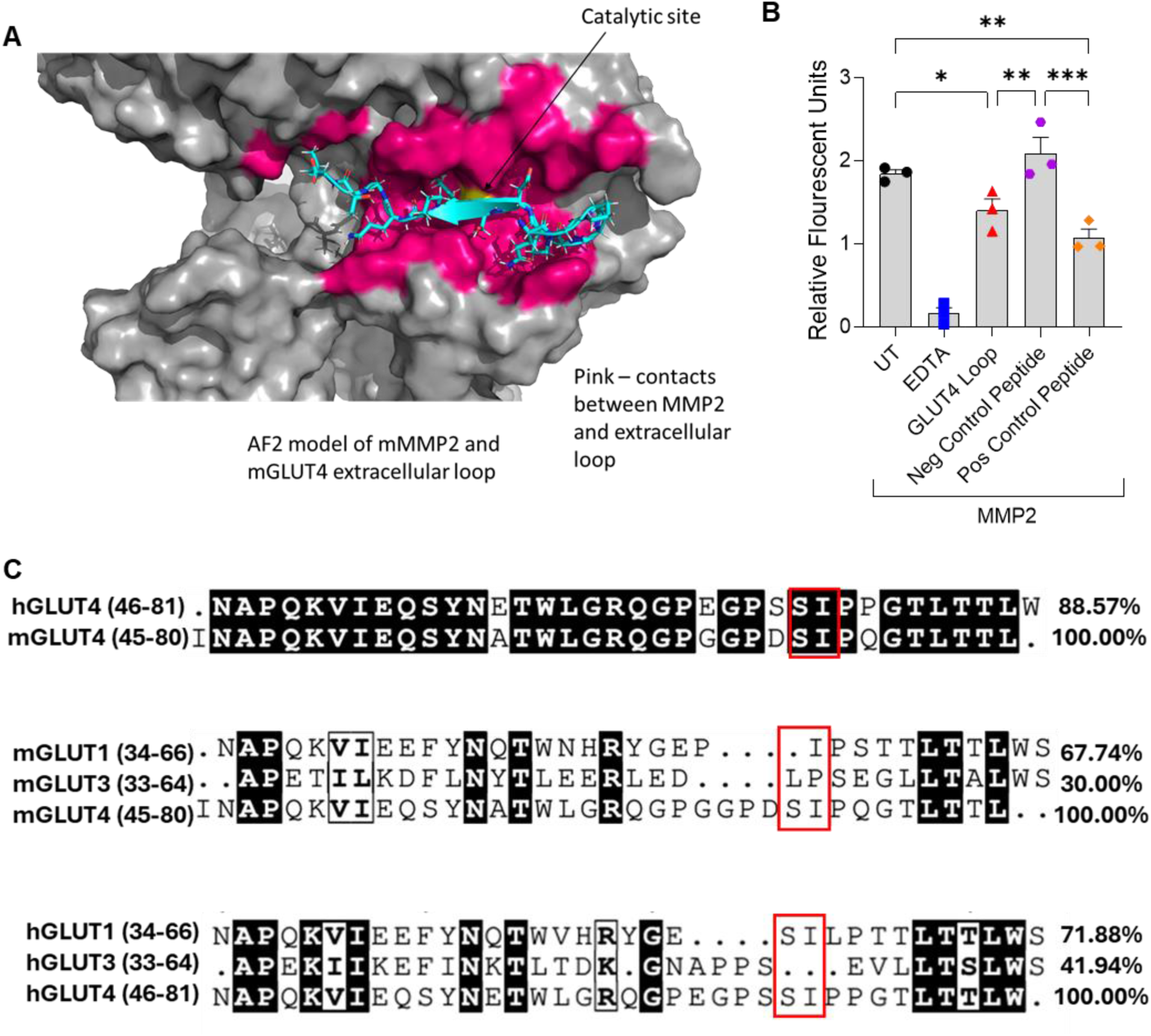
MMP2 cleavage site is localized in an extracellular domain of the GLUT4 receptor. **(A)** AlphaFold model of interaction between murine MMP2 and murine GLUT4 extracellular loop (residues 64-77). Pink represents contacts between MMP2 and the extracellular loop. **(B)** MMP fluorescent cleavage competition assay: MMP substrate, which becomes fluorescent after cleavage, was incubated at 37 °C with active MMP2, MMP2 with EDTA, MMP2 with GLUT4 loop peptide, MMP2 with negative control peptide, and MMP2 with positive control peptide. After 30 minutes, fluorescent intensity of the reactions was determined using a plate reader. **(C)** Sequence alignment of the extracellular loop between the first and second helix of human GLUT4 and murine GLUT4; murine GLUT1, murine GLUT3, and murine GLUT4; and human GLUT1, human GLUT3, and human GLUT4. Red box indicates MMP2 cleavage site. Values are expressed as means + SEM. *, p<0.05; **, p<0.01; ***, p>0.001 by nonparametric t-test (U-test). n=3 **(B)**

Sequence alignment between murine and human GLUT4 shows a high identity between the extracellular loops of both proteins (**Fig. 3C**), suggesting that the MMP2 cleavage of GLUT4 may be conserved among mice and humans. Furthermore, the sequence alignment of murine and human GLUT1 and GLUT3 to GLUT4 showed low identity (**Fig. 3C**), especially surrounding the proposed GLUT4 cleavage site, indicating that the cleavage event may be specific to GLUT4. Altogether, these results suggest that mouse GLUT4 has a potential MMP2 binding site, that is conserved in human GLUT4.

## DISCUSSION

Here, we show that MMP expression is increased in the gonadal WAT of IR mice, and the increased MMP expression is concentrated around CLSs. Furthermore, MMP2 is the highly active gelatinase in the gonadal WAT of HFD-fed obese mice. MMP2 treatment impaired glucose uptake and aerobic glycolysis in 3T3-L1 adipocytes, and inhibition of MMP2 activity reversed the impairment in metabolism. *In silico* structural analysis identified an MMP2 cleavage site on an extracellular domain of both murine and human GLUT4, and the GLUT4 extracellular domain was confirmed as a substrate for MMP2.

Apart from targeting extracellular matrix proteins, MMP2 affects cellular functions, such as, processing of the chemokine stromal cell-derived factor-α that causes neuronal apoptosis and neurodegeneration (13). MMP2 can be activated by the disruption of the cysteine-Zn^2+^ bond, either by proteolysis to an enzymatically active form (14) or by conformational changes caused by reactive oxygen-nitrogen species (RONS) (15). Proteolysis of MMP2 is facilitated by the formation of a trimeric complex of MMP2, tissue inhibitors of metalloproteinase 2 (TIMP2), and membrane-type 1 MMP (MT1-MMP or MMP-14) on the cell surface (16). Both TIMP2 and MT1-MMP are upregulated in obese adipose tissue, and adipocyte-specific overexpression of MT1-MMP in obese mice impairs lipid metabolism and insulin resistance (17; 18). Furthermore, RONS are increased in obese adipose tissue and produced by pro-inflammatory macrophages in CLSs (19; 20). Here we demonstrate increased MMP activity and MMP2 expression in CLSs, suggesting that RONS produced by the pro-inflammatory macrophages may be one of the primary activators of MMP2 during obesity.

MMP2 has also been identified as a regulator of adipocyte differentiation. Treatment of 3T3-L1 fibroblasts with a specific MMP2 inhibitor impairs adipogenic differentiation, primarily in the early stages of differentiation (21). Genetically *Mmp2* deficient mice are leaner and exhibit lower weight and growth rates at birth (22). Furthermore, *Mmp2* deficiency causes less fat accumulation in mice when challenged with an HFD (23). It remains unclear through what mechanism MMP2 promotes adipocyte differentiation. Due to MMP2’s effect on adipocyte differentiation, our studies utilized fully differentiated 3T3-L1 adipocytes before treatment with MMP2.

Our studies focus on the effect of MMP2 on glucose utilization and adipocyte metabolism using 3T3-L1 adipocytes as an *in vitro* model of white adipocytes. As *in vitro* models cannot fully recapitulate the complexities of *in vivo* physiology, future studies on mouse models are required to fully delineate the effect of MMP2 on adipocyte glucose metabolism during obesity. As genetic deletion of *Mmp2* results in mice with low body weight and defects in growth rate, utilization of a global inducible *Mmp2*-floxed mouse model is necessary to determine the physiological role of MMP2 on the development of insulin resistance and hyperglycemia.

Altogether, our study shows a novel function of MMP2, which may contribute to hyperglycemia during obesity. Furthermore, GLUT4 is expressed in aortic smooth muscle cells and reduction of GLUT4 levels in the aorta has been linked to hypertension and aortic dissection during metabolic disease (24; 25). This suggests that MMP2 may be a therapeutic target of interest to enhance glucose utilization in patients with a variety of metabolic diseases.

## AUTHOR CONTRIBUTIONS

M.D.L., R.J.G., and A.K.M. designed research; M.D.L., R.J.G., and A.K.M. performed research; B.L.G. and A.K.M. contributed new reagents/analytic tools; M.D.L., T.E.H., R.J.G., and A.K.M. analyzed data; and M.D.L. and A.K.M. wrote the paper.

## ACKNOWLEDGMENTS

The authors declare no conflicts of interest. We thank Jerald Whittley for technical assistance, East Carolina University Flow Core Facility, Research Histology Core, and Imaging Core for technical support. This project was supported by the Brody School of Medicine at ECU’s Mass Spectrometry Core Facilities which has received support from the Golden Leaf Foundation and from federal COVID-19 relief funds appropriated to ECU in North Carolina SL 2020-4.

## REFERENCES

1. Abel ED, Peroni O, Kim JK, Kim Y-B, Boss O, Hadro E, Minnemann T, Shulman GI, Kahn BB: Adipose-selective targeting of the GLUT4 gene impairs insulin action in muscle and liver. Nature 2001;409:729–733

2. Albers PH, Bojsen-Møller KN, Dirksen C, Serup AK, Kristensen DE, Frystyk J, Clausen TR, Kiens B, Richter EA, Madsbad S, Wojtaszewski JFP: Enhanced insulin signaling in human skeletal muscle and adipose tissue following gastric bypass surgery. American Journal of Physiology-Regulatory, Integrative and Comparative Physiology 2015;309:R510–R524

3. Pope BD, Warren CR, Parker KK, Cowan CA: Microenvironmental Control of Adipocyte Fate and Function. Trends in Cell Biology 2016;26:745–755

4. Lempicki MD, Gray JA, Abuna G, Murata RM, Divanovic S, McNamara CA, Meher AK: BAFF neutralization impairs the autoantibody-mediated clearance of dead adipocytes and aggravates obesity-induced insulin resistance. Front Immunol 2024;15:1436900

5. Lempicki MD, Paul S, Serbulea V, Upchurch CM, Sahu S, Gray JA, Ailawadi G, Garcia BL, McNamara CA, Leitinger N, Meher AK: BAFF antagonism via the BAFF receptor 3 binding site attenuates BAFF 60-mer-induced classical NF-kappaB signaling and metabolic reprogramming of B cells. Cell Immunol 2022;381:104603

6. Lumeng CN, Deyoung SM, Bodzin JL, Saltiel AR: Increased Inflammatory Properties of Adipose Tissue Macrophages Recruited During Diet-Induced Obesity. Diabetes 2007;56:16–23

7. Boumiza S, Chahed K, Tabka Z, Jacob M-P, Norel X, Ozen G: MMPs and TIMPs levels are correlated with anthropometric parameters, blood pressure, and endothelial function in obesity. Scientific Reports 2021;11

8. Hill DA, Lim H-W, Kim YH, Ho WY, Foong YH, Nelson VL, Nguyen HCB, Chegireddy K, Kim J, Habertheuer A, Vallabhajosyula P, Kambayashi T, Won K-J, Lazar MA: Distinct macrophage populations direct inflammatory versus physiological changes in adipose tissue. Proceedings of the National Academy of Sciences 2018;115:E5096–E5105

9. Huang W-C, Sala-Newby GB, Susana A, Johnson JL, Newby AC: Classical Macrophage Activation Up-Regulates Several Matrix Metalloproteinases through Mitogen Activated Protein Kinases and Nuclear Factor-κB. PLOS ONE 2012;7:e42507

10. Lee YJ, Heo YS, Park HS, Lee SH, Lee SK, Jang YJ: Serum SPARC and matrix metalloproteinase-2 and metalloproteinase-9 concentrations after bariatric surgery in obese adults. Obes Surg 2014;24:604–610

11. Yang J, Holman GD: Comparison of GLUT4 and GLUT1 subcellular trafficking in basal and insulin-stimulated 3T3-L1 cells. Journal of Biological Chemistry 1993;268:4600–4603

12. Chen EI, Kridel SJ, Howard EW, Li W, Godzik A, Smith JW: A unique substrate recognition profile for matrix metalloproteinase-2. J Biol Chem 2002;277:4485–4491

13. Zhang K, McQuibban GA, Silva C, Butler GS, Johnston JB, Holden J, Clark-Lewis I, Overall CM, Power C: HIV-induced metalloproteinase processing of the chemokine stromal cell derived factor-1 causes neurodegeneration. Nat Neurosci 2003;6:1064–1071

14. Cao J, Sato H, Takino T, Seiki M: The C-terminal Region of Membrane Type Matrix Metalloproteinase Is a Functional Transmembrane Domain Required for Pro-gelatinase A Activation. Journal of Biological Chemistry 1995;270:801–805

15. Viappiani S, Nicolescu AC, Holt A, Sawicki G, Crawford BD, León H, van Mulligen T, Schulz R: Activation and modulation of 72kDa matrix metalloproteinase-2 by peroxynitrite and glutathione. Biochemical Pharmacology 2009;77:826–834

16. Nagase H: Cell surface activation of progelatinase A (proMMP-2) and cell migration. Cell Research 1998;8:179–186

17. Nonino CB, Noronha NY, D. Araújo Ferreira-Julio M, Moriguchi Watanabe L, Cassia KF, Ferreira Nicoletti C, Rossi Welendorf C, Salgado Junior W, Rossi Silva Souza D, Augusta De Souza Pinhel M: Differential Expression of MMP2 and TIMP2 in Peripheral Blood Mononuclear Cells After Roux-en-Y Gastric Bypass. Frontiers in Nutrition 2021;8

18. Li X, Zhao Y, Chen C, Yang L, Lee H-H, Wang Z, Zhang N, Kolonin MG, An Z, Ge X, Scherer PE, Sun K: Critical Role of Matrix Metalloproteinase 14 in Adipose Tissue Remodeling during Obesity. Molecular and Cellular Biology 2020;40:1–24

19. Politis-Barber V, Brunetta HS, Paglialunga S, Petrick HL, Holloway GP: Long-term, high-fat feeding exacerbates short-term increases in adipose mitochondrial reactive oxygen species, without impairing mitochondrial respiration. American Journal of Physiology-Endocrinology and Metabolism 2020;319:E376–E387

20. Nakajima S, Koh V, Kua L-F, So J, Davide L, Lim KS, Petersen SH, Yong W-P, Shabbir A, Kono K: Accumulation of CD11c+CD163+ Adipose Tissue Macrophages through Upregulation of Intracellular 11β-HSD1 in Human Obesity. The Journal of Immunology 2016;197:3735–3745

21. Croissandeau G, CHrétien M, Mbikay M: Involvement of matrix metalloproteinases in the adipose conversion of 3T3-L1 preadipocytes. Biochemical Journal 2002;364:739–746

22. Itoh T, Ikeda T, Gomi H, Nakao S, Suzuki T, Itohara S: Unaltered Secretion of β-Amyloid Precursor Protein in Gelatinase A (Matrix Metalloproteinase 2)-deficient Mice. Journal of Biological Chemistry 1997;272:22389–22392

23. Van Hul M, Lijnen HR: A functional role of gelatinase A in the development of nutritionally induced obesity in mice. Journal of Thrombosis and Haemostasis 2008;6:1198–1206

24. Zheng H, Qiu Z, Chai T, He J, Zhang Y, Wang C, Ye J, Wu X, Li Y, Zhang L, Chen L: Insulin Resistance Promotes the Formation of Aortic Dissection by Inducing the Phenotypic Switch of Vascular Smooth Muscle Cells. Front Cardiovasc Med 2021;8:732122

25. Park JL, Loberg RD, Duquaine D, Zhang H, Deo BK, Ardanaz N, Coyle J, Atkins KB, Schin M, Charron MJ, Kumagai AK, Pagano PJ, Brosius FC, 3rd: GLUT4 facilitative glucose transporter specifically and differentially contributes to agonist-induced vascular reactivity in mouse aorta. Arterioscler Thromb Vasc Biol 2005;25:1596–1602

