## Supplementary material for "Matrix Metalloproteinase-2 as a novel regulator of glucose utilization by adipocytes": Online Supplemental Materials

### SUPPLEMENTAL MATERIALS AND METHODS

#### Isolation of SVF from Adipose Tissue

The left and the right gonadal white adipose tissue (WAT) was excised from the mouse and one of the fat pads was used for the isolation of the stromal vascular fraction (SVF). The tissue was minced with scissors and digested for 1 hour in Hanks Balanced Salt Solution (HBSS, Gibco) with 2% bovine serum albumin (BSA) and 2 mg/ml of collagenase type I shaking at 37 °C. After digestion, tissue was filtered through a 500 μM nylon mesh strainer and washed with HBSS with 2% BSA. The cells were centrifuged (450xg for 5 min) to allow the stromal vascular fraction (SVF) to separate with the adipocytes floating on the top of the buffer. The SVF pellet was collected, and red blood cells were lysed. Cells were washed, centrifuged, and resuspended in various buffers depending on downstream use.

#### Flow Cytometry

The SVF from WAT was isolated as described above. The cells were blocked with purified anti-mouse CD16/32 (BioLegend, San Diego, CA; #101302) and stained with the following fluorescent conjugated antibodies from BioLegends and FisherScientific for 30 minutes on ice: PE Cyanine7 anti-F4/80 (#25-4801-82), Brilliant Blue 515 anti-CD45 (#564590), and APC anti-CD11b (#17-0112-83), Brilliant Violet 711 anti-CD206 (#141727), and Brilliant Violet 605 anti-CD86 (#105037). Invitrogen DAPI (4',6-diamidino-2-phenylindole, dihydrochloride) (#D1306) was used for staining the dead cell. The samples were run on the flow cytometer machine Cytek Biosciences Cytek Aurora, USA. Data analysis and quantification were performed using FlowJo v10.6.2.

**Plasma insulin quantification**

Fasting plasma insulin levels were measured with an Ultra-Sensitive Mouse Insulin ELISA Kit (Crystal Chem, Elk Grove Village, IL; # 90080) per manufacturer instructions.

**Western Blot**

Adipose tissue was homogenized in a bead mill with TPER buffer (ThermoFisher Scientific #78510) containing a protease inhibitor cocktail (Sigma-Aldrich). The extracted protein was analyzed using 4-12% Tris-Glycine SDS-PAGE. The primary antibodies used are anti-AKT and anti-pAKT and the IRDye-conjugated secondary antibodies were from LI-COR. The blots were detected, and band intensities were quantified using LI-COR's Image Studio software version 5.2.5.

**Real-Time qPCR**

SVF was isolated as described above and resuspended in TRIzol reagent (ThermoFisher, # 15596026) and RNA fraction was isolated per manufacturer's instructions. RNA was purified with RNeasy Mini Kit (Qiagen, # 74104) per manufacturer's instructions. cDNA was synthesized and real-time PCR was performed on a CFX Connect™ Real-Time PCR Detection System as described previously (1; 2). The abundance of *B2m* mRNA was used for relative quantification of *Tnfα* mRNAs in the SVF.

**Immunohistochemistry**

Five micrometer gonadal WAT cross-sections were hydrated with HistoClear Clearing Agent (MilliporeSigma), 100% ethanol, 90% ethanol, 70% ethanol, 20% ethanol, and then water. Antigen retrieval was performed by boiling the slides in the Antigen Unmasking Solution (Vector

Laboratories, Burlingame, CA; #H-3300). After antigen retrieval, the sections were treated with 10% donkey serum and incubated with primary Ab for 14 to 16 hours at 4°C. Primary antibodies were detected by fluorescent secondary antibodies. The slides were mounted with ProLong Gold Antifade Mountant with DAPI (Invitrogen, #P36931) before image acquisition on a Laxco LMI 6000 Series Inverted microscope (Fisher Scientific) or EVOS M5000 (ThermoFisher). The primary antibodies used were anti-FABP4/A-FABP (Novus Biologics; #AF1443), anti-Mac-2 (Cederlane; #CL8942AP), anti-MMP2 (ThermoFisher, #10373-AP). The secondary antibodies used are Donkey anti-Goat Alexa Fluor Plus 488 (ThermoFisher; #A32814), Donkey anti-Rabbit Alexa Fluor Plus 555 (ThermoFisher; #A32794), and Donkey anti-Rat Alexa Fluor Plus 647 (ThermoFisher; A48272).

**Gelatin Zymogram**

Adipose tissue was homogenized in a bead mill with TPER buffer (ThermoFisher Scientific #78510) containing a protease inhibitor cocktail (Sigma-Aldrich). Protein concentration was determined by Pierce BCA protein assay (ThermoFisher Scientific, #23227). Non-reduced total lysate was loaded at a concentration of 15 μg/mL onto a 12% SDS-polyacrylamide gel containing 1 mg/mL of porcine gelatin (Sigma Aldrich, #G1890). Following electrophoresis, the gels were washed and renatured with washing buffer (2.5% Triton X-100, 50 mM Tris-HCl pH7.5, 5 mM $CaCl_2$, 1 μM $ZnCl_2$) with two 30min washes at room temperature. Gels were then incubated with substrate buffer (1% Triton X-100, 50 mM Tris-HCl pH 7.5, 5 mM $CaCl_2$, 1 μM $ZnCl_2$) at 37°C for 24 hrs. Gels were stained with Coomassie Blue staining solution (40% Methanol, 10% Acetic Acid, 0.5% Coomassie Blue) for 30 min at room temperature and then destained with destaining solution (40% Methanol, 10% Acetic Acid) for 1hr.

**Preparation of Bone-Marrow-Derived Macrophages and Differentiation of 3T3-L1 Adipocytes**

The bone-marrow-derived macrophages and 3T3-L1 adipocytes were prepared as described before (1). Briefly, bone marrow cells from the femur and tibia of WT C57BL6/J mice were collected, red blood cells were lysed with lysis buffer and the cells were cultured at a density of 1 to $2\times10^6$ cells/mL in RPMI 1640 medium containing 10% heat-inactivated fetal bovine serum (HI FBS) and 1× antibiotic-antimycotic, supplemented with 20% L929 (NCTC clone 929; ATCC, Manassas, VA) culture supernatant. The culture medium was changed on the fourth day and then on every second day. M1 or M2 macrophages were induced by incubation with 1μg/mL of LPS and 50 ng/mL of IFNγ or 20 ng/mL of IL-4 respectively overnight at 37°C.

3T3-L1 fibroblasts were cultured in DMEM supplemented with 10% heat-inactivated (HI) FBS at 37°C, 95% relative humidity, and 5% CO2. For differentiation into adipocytes, cells were grown to confluence and treated for four days with differentiation media containing DMEM, 10% HI FBS, 1x antibiotic-antimycotic, 0.25 U/mL insulin, 0.5 mM 3-isobutyl-1-methylxanthine, 0.025 mM dexamethasone, on day 5 cells were treated with post-differentiation media containing DMEM, 10% HI FBS, 0.25 U/mL insulin and then adipocytes were maintained in DMEM with 10% heat-inactivated FBS. 3T3-L1 adipocytes were used in glucose uptake and seahorse experiments on day 6 of differentiation.

**Glucose Uptake**

3T3-L1 adipocytes, 5 days after differentiation, were seeded one day prior to glucose uptake assay on 24-well plates coated with Poly-L-lysine solution (P4832, Sigma-Aldrich, USA) and incubated overnight at 37°C to adhere. Cells were washed with phosphate buffered saline (PBS) and incubated for 2 hours in low Glucose DMEM (5 mM glucose). For the MMP2 concentration titration assay, cells were left untreated or treated with 200 nM insulin plus 200, 400, or 800 ng/mL of activated MMP2 (Biolegend, #55402) in low Glucose DMEM for 20 min at 37 °C. For the MMP2 inhibitor assay, cells were left untreated or treated with 200 nM insulin with 400 ng/mL active MMP2, 1 μM MMP2 inhibitor (Cayman chemicals, #19644), or active MMP2 with MMP2 inhibitor in low glucose DMEM for 20 min at 37°C. Cells were then treated with 200 μg/mL 2-NBDG (2-(N-(7-Nitrobenz-2-oxa-1,3-diazol-4-yl)Amino)-2-Deoxyglucose) (Thermofisher, #N13195) for 10 min at 37 °C. After incubation, cells were washed, removed with accutase (StemCell Technologies), and glucose uptake was analyzed by flow cytometer. Cells were treated with Cytochalasin D (10 μmol/L; Cayman Chemicals) after 1 hr of incubation with low glucose DMEM as a negative control for glucose uptake.

**Seahorse**

The 3T3-L1 fibroblasts were seeded on XFe24 well microplates coated with Poly-L-lysine solution (P4832, Sigma-Aldrich, USA) and differentiated into adipocytes as described earlier or seeded one day prior to the Seahorse experiment in coated microplates. A glycolysis stress test on Seahorse XFe24 extracellular flux analyzer was performed using a Seahorse XFe24 Extracellular Flux Analyzer as described previously (3). Concentrations of the drugs injected are: 1st injection D-(+)-Glucose (20 mM, Sigma-Aldrich, G7021), 2nd injection Oligomycin A (10 μM), and the 3rd injection 2-Deoxy-D-glucose (80 mM, Sigma-Aldrich, #D6134). Extracellular acidification rate (ECAR) was measured three times before the first injection and after each subsequent injection.

**MMP Substrate Cleavage Assay**

In black 96-well plate 5 nM (400 ng/mL ) of MMP2 (BioLegend, #554402), 1 μM of MMP2 inhibitor (Cayman Chemicals, #19644), 5 nM of MMP2 with 1 μM of MMP2 inhibitor, or 5 nM of MMP2 with 50 mM EDTA as negative control were incubated with 5 μM of 520 MMP FRET Substrate (Anaspec, #AS-60568-01) for 30 minutes at 37 °C in MMP substrate buffer (50 mM Tris-HCl pH 7.5, 10mM $CaCL_2$, 150 mM NaCl, 5 μM $ZnCl_2$). Fluorescence was measured on Varioskan LUX (ThermoFisher) at 494/521 excitation/emission.

For the substrate competition assay, 5 nM of MMP2 with 500 μM of GLUT4 loop peptide (RQGPGGPDSIPQGTL, GenScript), 500 μM of negative control peptide (LTGQPISDPGGPGQR, GenScript), 500 μM positive control peptide (VPLSLYSG, Bachm, #4109381), or 50 mM EDTA were incubated with 5 μM of 520 MMP FRET Substrate (AnaSpec) for 30 minutes in MMP substrate buffer.

**Statistical Analysis**

The data were analyzed by using GraphPad Prism 8 (GraphPad Software, La Jolla, CA) and Excel (Microsoft Corporation, Redmond, WA), and they are presented as means ± SEM. Differences between the mean values of the two groups were determined by using t-tests. The means of multiple groups were compared by using a one-way analysis of variance. The D'Agostino-Pearson normality test was performed on each group. If the P value was not significant (>0.05), a two-tailed parametric test was used; if the P value was significant (<0.05) or if the number of samples was four, a two-tailed nonparametric t-test (U-test) was used to determine significant differences between the groups. In the multiple comparisons, if a significant difference was found among the groups, pairs of groups were compared by using a parametric or nonparametric t-test. Statistical analyses are provided in each figure legend.

Differences between the groups were considered significant when $P < 0.05$. P values $>0.05$ are indicated in the graphs.

**SUPPLEMENTAL FIGURES**

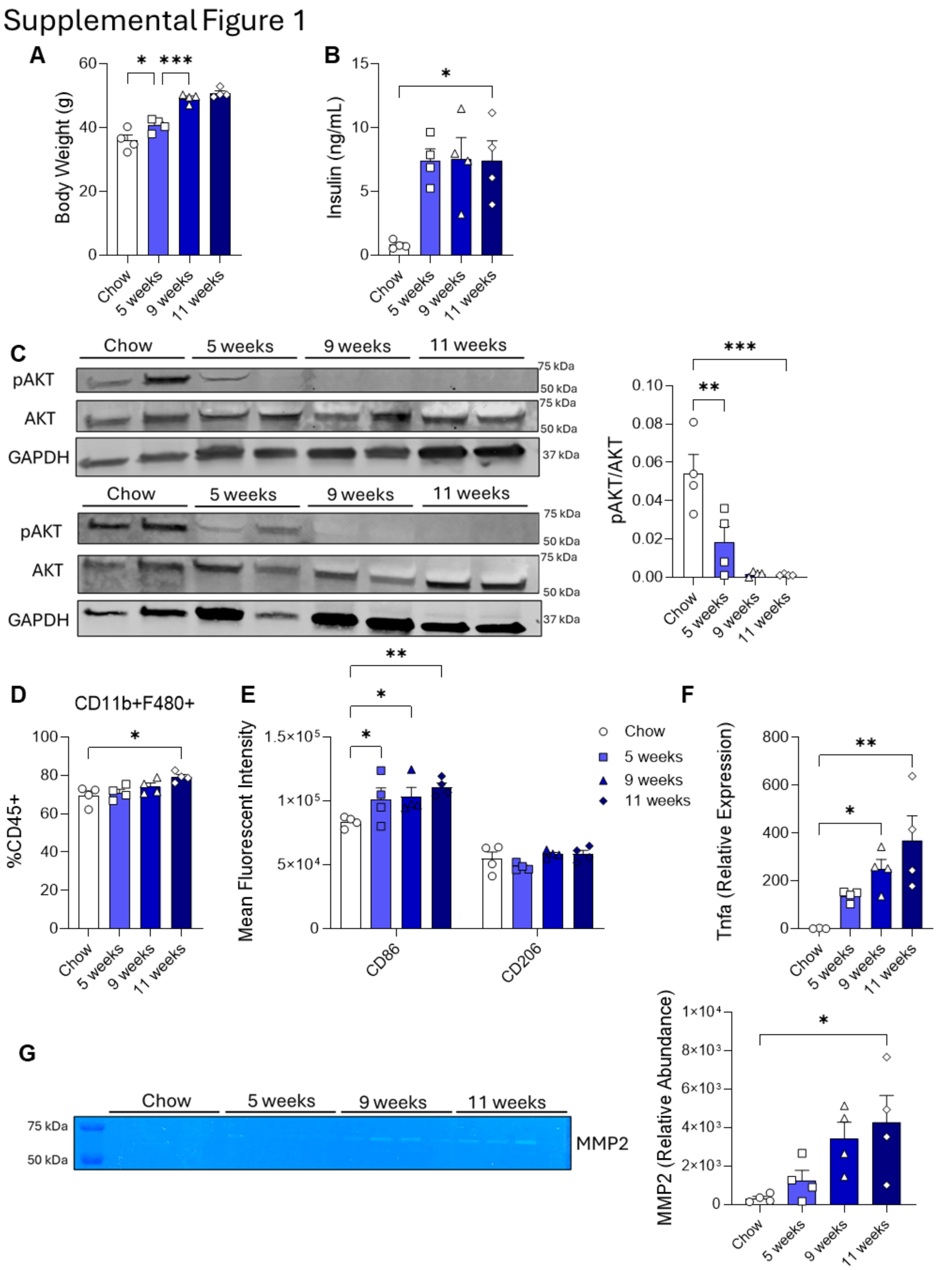

**Supplemental Figure 1. (Supplement to Figure 1) (A)** Body weight of mice, **(B)** Fasting plasma insulin levels, **(C)** Western blot and quantification of phosphorylated AKT, AKT, and GAPDH of gonadal WAT, **(D)** Percent population of total macrophages out of CD45+ cells in the SVF from gonadal WAT, **(E)** Mean fluorescent intensity of CD86 (M1 macrophage marker) and CD206 (M2 macrophage marker) on macrophages in SVF from gonadal WAT, **(F)** *Tnfα* gene expression in the SVF, and, **(G)** Gelatin zymogram of gonadal WAT lysate from mice on HFD for 5, 9, and 11 weeks or on NCD for 11 weeks. Values are expressed as means + SEM. *, $p<0.05$; **, $p<0.01$; ***, $p>0.001$ by nonparametric t-test (U-test). n=4 **(A-G).**

Supplemental Figure 2

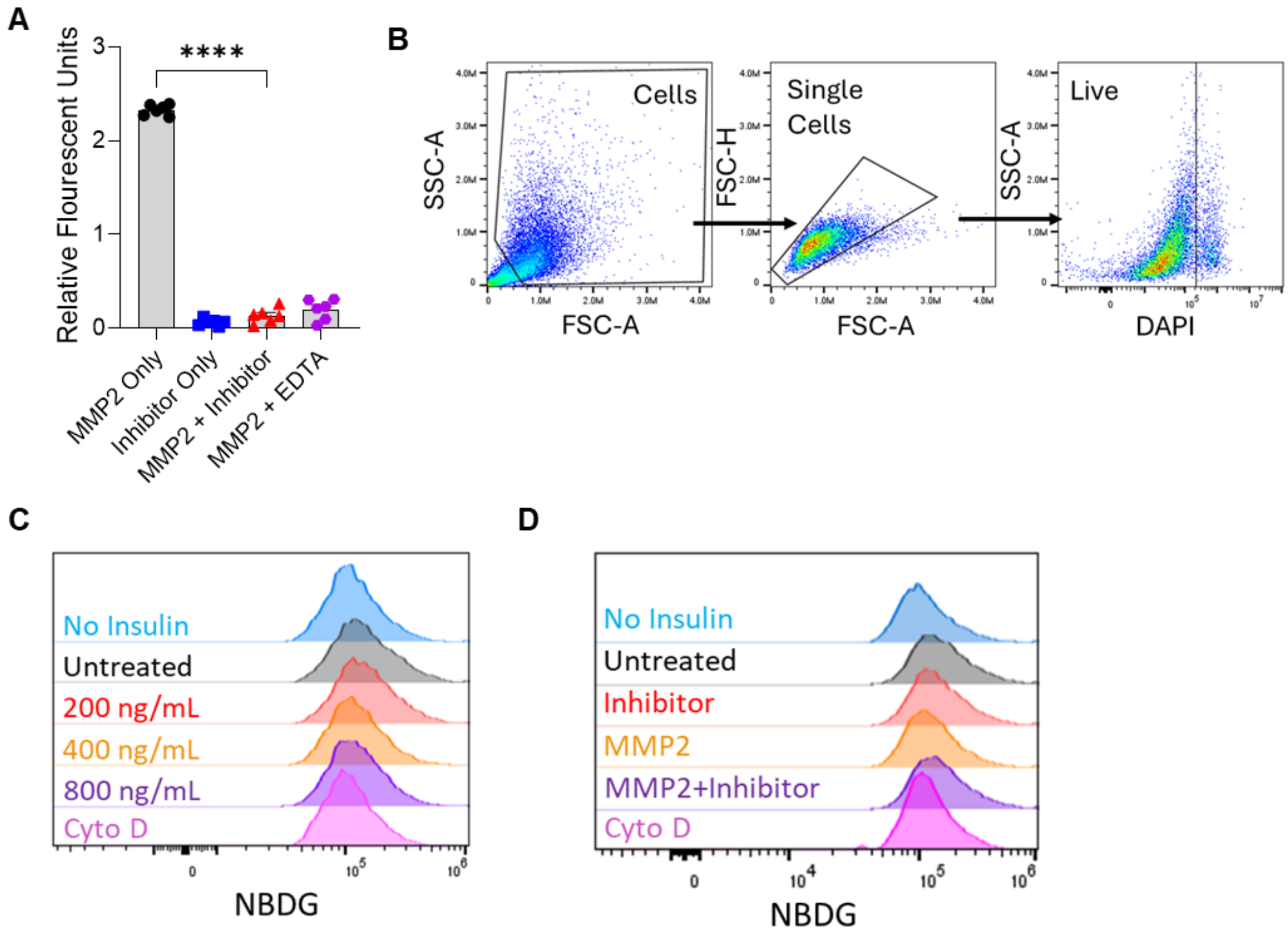


**Supplemental Figure 2. (Supplement to Figure 2). (A)** MMP fluorescent cleavage assay and validation of MMP2 inhibitor: MMP substrate, which becomes fluorescent after cleavage, incubated with active MMP2, MMP2 inhibitor only, MMP2 with MMP2 inhibitor, or MMP2 with EDTA (inhibits MMP activity). **(B)** Flow cytometry gating for adipocytes in glucose uptake assay. **(C)** Histograms of median fluorescent intensity (MFI) of 2-NBDG of adipocytes after treatment with 200 ng/mL, 400 ng/mL, or 800 ng/mL of active MMP2. **(D)** Histograms of median fluorescent intensity (MFI) of 2-NBDG of adipocytes after treatment with 400 ng/mL of active MMP2, MMP2 inhibitor, or MMP2 with MMP2 inhibitor. Values are expressed as means + SEM. ****, $p<0.0001$ by parametric unpaired t-test. n=6 **(A-D)**

### REFERENCES


1. Meher AK, Sharma PR, Lira VA, Yamamoto M, Kensler TW, Yan Z, Leitinger N: Nrf2 deficiency in myeloid cells is not sufficient to protect mice from high-fat diet-induced adipose tissue inflammation and insulin resistance. Free Radic Biol Med 2012;52:1708-1715

2. Kadl A, Meher AK, Sharma PR, Lee MY, Doran AC, Johnstone SR, Elliott MR, Gruber F, Han J, Chen W, Kensler T, Ravichandran KS, Isakson BE, Wamhoff BR, Leitinger N: Identification of a novel macrophage phenotype that develops in response to atherogenic phospholipids via Nrf2. Circ Res 2010;107:737-746

3. Lempicki MD, Paul S, Serbulea V, Upchurch CM, Sahu S, Gray JA, Ailawadi G, Garcia BL, McNamara CA, Leitinger N, Meher AK: BAFF antagonism via the BAFF receptor 3 binding site attenuates BAFF 60-mer-induced classical NF-kappaB signaling and metabolic reprogramming of B cells. Cell Immunol 2022;381:104603
